# A natural genetic variation screen identifies insulin signaling, neuronal communication, and innate immunity as modifiers of hyperglycemia in the absence of *Sirt1*

**DOI:** 10.1101/2021.09.13.460173

**Authors:** Katie G. Owings, Rebecca A.S. Palu

## Abstract

Variation in the onset, progression, and severity of symptoms associated with metabolic disorders such as diabetes impairs the diagnosis and treatment of at-risk patients. Diabetes symptoms, and patient variation in these symptoms, is attributed to a combination of genetic and environmental factors, but identifying the genes and pathways that modify diabetes in humans has proven difficult. A greater understanding of genetic modifiers and the ways in which they interact with metabolic pathways could improve the ability to predict a patient’s risk for severe symptoms, as well as enhance the development of individualized therapeutic approaches. In this study we use the *Drosophila* Genetic Reference Panel (DGRP) to identify genetic variation influencing hyperglycemia associated with loss of *Sirt1* function. Through analysis of individual candidate functions, physical interaction networks, and Gene Set Enrichment Analysis (GSEA) we identify not only modifiers involved in canonical glucose metabolism and insulin signaling, but also genes important for neuronal signaling and the innate immune response. Furthermore, reducing the expression of several of these candidates suppressed hyperglycemia, making them ideal candidate therapeutic targets. These analyses showcase the diverse processes contributing to glucose homeostasis and open up several avenues of future investigation.

## INTRODUCTION

Metabolic diseases, and in particular diabetes, are one of the most pressing health crises in the developed world, with incidences continuing to rise in the last 20 years (CDC 2020). 37.7% of adults in the US are diagnosed as obese and 10.5% as having some form of diabetes, and it is estimated that millions more go undiagnosed (Flegal *et al.* 2016; CDC 2020). What is more, the monetary cost of these disorders to the public has grown to astronomical levels. It is estimated that $237 billion in direct costs and at least $90 billion in indirect costs were spent on healthcare related to diabetes and its various complications in 2017 alone, up ∼25% from 2012 (Flegal *et al.* 2016). A focused effort has been made to understand both the genetic and environmental contributors to metabolic homeostasis, as well as to the disruption of that homeostasis that leads to disease (Barroso and McCarthy 2019).

Unfortunately, even identifying these contributors has proven difficult. Metabolic diseases are complex, and the onset, progression, and ultimately the severity of any individual case is dependent upon a myriad of genetic and environmental variables and the ways in which they interact with one another (Queitsch *et al.* 2012; Barroso and McCarthy 2019). Even when there is a strong familial link, phenotypic heterogeneity in disease phenotypes can make it difficult to identify at-risk patients or make accurate prognostic predictions (Udler *et al.* 2019). This is particularly true when it comes to predicting complications of diabetes such as neuropathy, retinopathy, or kidney disease (Barroso and McCarthy 2019; Cabrera *et al.* 2020). Much of this variation is due to inter-individual differences in genetic background, including silent cryptic genetic variation that is revealed upon disease or stress (Queitsch *et al.* 2012; Chow 2016; Barroso and McCarthy 2019).

One example of this kind of symptom heterogeneity can be observed in disease associated with the deacetylase *SIRT1*. This highly conserved gene was originally identified as a histone deacetylase important in heterochromatin formation in yeast (Shore *et al.* 1984; Ivy *et al.* 1986; Rine and Herskowitz 1987). Since then, SIRT1 and its paralogs (the sirtuins) have been found to have a number of additional targets, many of which are transcription factors and enzymes with key roles in metabolic homeostasis (Brunet *et al.* 2004; Picard *et al.* 2004; Rodgers and Puigserver 2007; Li *et al.* 2007; Yang *et al.* 2009; Palu and Thummel 2016). Importantly, as part of their enzymatic reaction, sirtuins consume the cofactor NAD, which also serves as an electron carrier in central metabolic pathways such as glycolysis and the TCA cycle. Sirtuin enzymatic activity, therefore, is directly linked to the availability of this cofactor and thus is responsive to the energetic state of the cell. This information is then conveyed to the targets, whose acetylation state alters their activity and stability in the cell (Nogueiras *et al.* 2012). With this centralized role in regulating the response of metabolic factors to cellular energy availability, it is unsurprising that variation in *SIRT1* has been linked to, among other things, the development of diabetes (Zillikens *et al.* 2009; Botden *et al.* 2012; Biason-Lauber *et al.* 2013; Zhao *et al.* 2017).

Elucidating the mechanism behind this link, however, has proven difficult. Loss-of-function and gain-of-function studies in model systems have demonstrated a clear role for SIRT1 in metabolic homeostasis, but the actual impacts on the animals in question have frequently been contradictory (Boutant and Cantó 2014). It is likely that at least some of these contradictory results stem from differences in genetic background between the animals used in the various studies. Understanding the role of this variation and the genes or pathways which modify metabolic disease will enable the development of improved diagnosis, prediction of prognosis, and personalized treatment strategies for patients.

Model organism tools, such as the *Drosophila* Genetic Reference Panel (DGRP), provide a way to study of the impact of natural genetic variation on diseases such as diabetes (Mackay *et al.* 2012; He *et al.* 2014; Ivanov *et al.* 2015; Nelson *et al.* 2016; Jehrke *et al.* 2018; Everman *et al.* 2019). The DGRP is a collection of ∼200 isogenic strains derived from a wild population, such that each strain represents one wild-derived genome (Mackay *et al.* 2012). The variation in the DGRP is well tolerated under healthy, non-disease conditions and allows for the identification of genetic polymorphisms that are associated with phenotypic variation in models of human disease (Chow and Reiter 2017). Importantly, the availability of full-genome sequence for these strains allows for genome-wide association analyses that link quantitative phenotypes with genetic variation and modifier genes.

The utility of the DGRP in identifying candidate modifiers of metabolic disease has already been demonstrated numerous times, with screens associated with misfolded insulin, high sugar and high fat feeding, and starvation resistance already documented (Mackay *et al.* 2012; He *et al.* 2014; Ivanov *et al.* 2015; Nelson *et al.* 2016; Jehrke *et al.* 2018; Everman *et al.* 2019). While some of these used biochemical assays to precisely measure metabolite levels in flies as a quantitative phenotype for the screen (Nelson *et al.* 2016; Jehrke *et al.* 2018; Everman *et al.* 2019), several used more general physiological measurements such as starvation resistance and lifespan (Mackay *et al.* 2012; Ivanov *et al.* 2015). Although effective, the use of this kind of general readout reduces the specificity of the modifiers identified. Many genetic factors impact survival and could lead to a high background signal. The same could be true for otherwise wild-type flies subjected to different environmental conditions, even when more precise assays are used as a quantitative phenotype (Nelson *et al.* 2016). A multitude of pathways and processes impact the response to dietary changes through feeding rate, hormone secretion, anabolism and catabolism rates, and nutrient absorption. Using a specific genetic model of disease and then focusing on a specific phenotype, such as hyperglycemia, may reduce some of the noise and increase specificity of the modifiers identified.

Furthermore, recent studies using the DGRP have demonstrated that the top candidate modifier genes and pathways differ when different but related models of genetic disease are screened (Chow *et al.* 2016; Palu *et al.* 2019). These results reinforce the idea that different causative genes and mutations will interact with different pathways over the course of disease. It also highlights the importance of exploring multiple disease models in something so diverse as diabetes, and the utility of the DGRP in precisely distinguishing modifiers of a particular genotype-phenotype combination.

In this study, we report the results of a natural variation screen in a model of *Sirt1* loss-of-function. Loss of this gene in *Drosophila* has been shown to lead to progressive metabolic dysfunction including obesity, hyperglycemia, and ultimately insulin resistance (Palu and Thummel 2016). Our study design is specifically focused on the phenotype of hyperglycemia when *Sirt1* expression is disrupted using RNAi in the adipose and liver-like fat body organ. We observed substantial phenotypic variation across the DGRP for hyperglycemia associated with loss of *Sirt1*. Using genome-wide association analysis, pathway enrichment, and the generation of a physical interaction network, we identified a number of modifying pathways and processes, several of which have known roles in central carbon metabolism, the immune response, and the kind of neuronal signaling and communication expected to influence the neuroendocrine cells responsible for insulin secretion in *Drosophila*. Finally, we confirmed that loss of several of the top candidate modifier genes significantly alters glucose levels in the *Sirt1* RNAi model. Our findings highlight exciting new areas of study for modifiers of *Sirt1* function, glucose homeostasis, and insulin sensitivity.

## METHODS

### Fly stocks and maintenance

Flies were raised at room temperature on a diet based on the Bloomington Stock Center standard medium with malt. Experimental crosses were maintained on a media containing 6% yeast, 6% dextrose, 3% sucrose, and 1% agar, with 0.6% propionic acid and 0.1% p-Hydroxy-benzoic acid methyl ester in 95% ethanol included as antifungal agents. Flies subjected to an overnight fast were transferred to media containing only 1% agar in water. The *R4*>*Sirt1i* strain, which serves as the model of hyperglycemia in this study, is derived from an *R4*-*GAL4* strain (BDSC 33832) outcrossed to *w^1118^* (Palu and Thummel 2016) and a *Sirt1* RNAi strain from the Bloomington Stock Center (32481). 185 strains from the DGRP were used for the modifier screen (Tables S1-S3), wherein virgin females carrying the *R4*>*Sirt1i* model were crossed to males of the DGRP strains. Male F1 progeny carrying *R4*>*Sirt1i* were separated and aged for one to two weeks. These flies were then either collected under ad libitum fed conditions or fasted overnight and then collected. The following RNAi and control strains are from the Bloomington Stock Center: *CG4168* RNAi (28636), *CG5888* RNAi (62175), *uif* RNAi (38354), *CTPSyn* RNAi (31924), *smt3* RNAi (36125), *ilp5* RNAi (33683), *Vha55* RNAi (40884), *snRNP*-*U1*-*70k* RNAi (33396), *CG10265* RNAi (43294), *CG15803* RNAi (51449), *Roe* RNAi (57836), *CG34353* RNAi (58291), *CadN2* RNAi (27508), *Ace* RNAi (25958), *CG43897* (31560), *dsxc73A* RNAi (56987), *bgm* RNAi (56979), *CG3407* RNAi (57762), control *attP40* (36304), and control *attP2* (36303).

### Glucose Assay

Samples of five flies each were collected at one or two weeks of age and washed in 1×PBS. Samples were then either frozen in liquid nitrogen and stored long-term at −80C or immediately homogenized in 100 uL 1× PBS. Frozen samples were kept frozen until immediately upon addition of PBS and homogenization. After homogenization samples were subjected to heat inactivation of enzymes at 70C for approximately 10 minutes. Glucose was measured undiluted from the lysate using the Sigma HK Glucose Assay kit as described (Tennessen *et al.* 2014).

### Protein Assay

Prior to heat inactivation, 10 uL of the fly lysate isolated for glucose measurement was saved and kept on ice. Protein samples could then be stored long term at −80°C. Samples were centrifuged for up to 5 minutes at room temperature and protein measured from the supernatant after a 1:10 dilution using the Sigma Protein Assay Reagent as described (Tennessen *et al.* 2014).

### Phenotypic analysis and genome-wide association

For each DGRP line, glucose was measured from 3 samples of 5 flies each. The P-values for association of genetic background and glucose concentration were calculated using one-way ANOVA on R software taking into account all collected data points for each experiment. Average glucose concentration was used for the genome-wide association (GWA). GWA was performed as previously described (Chow *et al.* 2016; Palu *et al.* 2019). DGRP genotypes were downloaded from the website, http://dgrp.gnets.ncsu.edu/. Non-biallelic sites were removed. A total of 3,636,891 variants were included in the analysis. Mean eye glucose concentration for 555 samples representing 2775 DGRP/R4>Sirt1i F1 progeny were regressed on each SNP. To account for cryptic relatedness (He *et al.* 2014; Huang *et al.* 2014), GEMMA (v. 0.94) (Zhou and Stephens 2012) was used to both estimate a centered genetic relatedness matrix and perform association tests using the following linear mixed model (LMM):

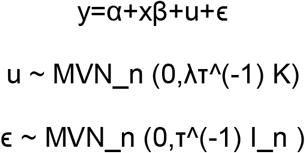

where, as described and adapted from Zhou and Stephens 2012, y is the n-vector of average glucose concentration for the n lines, α is the intercept, x is the n-vector of marker genotypes, β is the effect size of the marker. u is a n × n matrix of random effects with a multivariate normal distribution (MVN_n) that depends on λ, the ratio between the two variance components, ⊺^(−1), the variance of residuals errors, and where the covariance matrix is informed by K, the calculated n × n marker-based relatedness matrix. K accounts for all pairwise non-random sharing of genetic material among lines. ϵ, is a n-vector of residual errors, with a multivariate normal distribution that depends on ⊺^(−1) and I_n, the identity matrix. Quantile-quantile (qq) plots demonstrate an appropriate fit to the LMM at the positive end of the plot, but a greater number of points than expected by chance with an insignificant p-value (Figure S1). Genes were identified from SNP coordinates using the BDGP R54/dm3 genome build. A SNP was assigned to a gene if it was +/− 1 kb from a gene body.

### RNAi Validation

Virgin females from the *R4*>*Sirt1i* model were crossed to males carrying RNAi constructs targeting candidate modifiers of those models, and the glucose levels of F1 male progeny expressing both *Sirt1i* and the modifier RNAi construct specifically in the fat body were measured as described above on 4-5 samples of 5 male flies each. Glucose concentrations from RNAi-carrying strains are compared directly to genetically matched *attP40* or *attP2* controls using a Dunnett’s multiple comparisons test. Glucose measurements are normalized to the appropriate genetically matched controls. Normalized controls from individual experiments are compared in Figure S2. Standard deviation did not significantly vary between controls for individual experiments.

### Bioinformatics Analysis

Genetic polymorphisms were associated with candidate genes within 1 kb of the polymorphism. Information about candidate genes and their human orthologues was gathered from a number of databases including Flymine, Flybase, OMIM, and NCBI. Physical interaction maps were generated using the GeneMANIA plugin on Cytoscape (version 3.8.2) (Shannon *et al.* 2003; Montojo *et al.* 2010). GSEA was run to generate a rank-list of genes based on their enrichment for significantly associated polymorphisms. For GSEA analysis, polymorphisms within 1kb of more than 1 gene were assigned to one gene based on a priority list of exon, UTR, intron, and upstream or downstream. Genes were assigned to GO categories, and calculation of enrichment score was performed as described (Subramanian *et al.* 2005). Categories with ES scores > 0 (enriched for associated genes with low p-values), gene number > 3, and p-values <0.05 were included in the final output.

## RESULTS

### Glucose levels in R4>Sirt1i flies vary with genetic background in a consistent pattern across multiple conditions

Loss of *Sirt1* expression leads to progressive hyperglycemia, obesity, and insulin resistance. To model the hyperglycemia that is commonly associated with diabetes, we reduced the expression of the deacetylase *Sirt1* specifically in the fat body of *Drosophila melanogaster*. RNAi targeting *Sirt1* specifically in the fat body reproduces the hyperglycemia phenotype, with an approximately 50-60% increase in whole fly glucose levels (P = 0.02, Figure S3) (Palu and Thummel 2016). The fat body in *Drosophila* performs functions normally undertaken by the adipose and liver tissues in humans (Géminard *et al.* 2009; DiAngelo and Birnbaum 2009; Arrese and Soulages 2010). While there are likely roles for *Sirt1* in other metabolic tissues, its function in the fat body clearly contributes to the maintenance of glucose homeostasis and insulin sensitivity over time (Palu and Thummel 2016). Modifiers of insulin signaling or glucose metabolism could alter the degree of hyperglycemia in this model.

The degree of hyperglycemia is determined by a biochemical assay that measures the concentration of glucose in a whole-fly lysate (Tennessen *et al.* 2014). This is a quantitative assay where higher concentrations of glucose correlate with more severe hyperglycemia, and lower concentrations of glucose correlate with milder disease, or potentially hypoglycemia. Glucose was measured specifically in males, as adult female *Drosophila* devote much of their physiological output to egg production (Millington and Rideout 2018). Previous studies have demonstrated that adult males provide a consistent model for metabolic homeostasis in the fly (Sieber and Thummel 2009; Tennessen *et al.* 2014; Palu and Thummel 2016; Barry and Thummel 2016; Beebe *et al.* 2020).

Loss of *Sirt1* is induced using the *GAL4*/*UAS* system, where *R4*-*GAL4* drives expression of *UAS*-*Sirt1* RNAi (Figure S4A). *R4*-*GAL4* is strongly expressed primarily in the fat body of fly, starting in early development and continuing through adulthood (Lee and Park 2004). The line containing this model (*R4*>*Sirt1i*) serves as the donor strain that was crossed to each DGRP strain. Females from the donor strain were crossed with males of each of 185 DGRP strains to generate F1 progeny lacking *Sirt1* expression in the fat body. The progeny received 50% of their genome from the maternal donor strain and 50% from the paternal DGRP strain (Figure S4B). Therefore, we are measuring the dominant effect of the DGRP background on the *Sirt1* RNAi hyperglycemia phenotype. This experimental design is similar to a model of *NGLY1* deficiency using RNAi that was also crossed to the DGRP (Talsness *et al.* 2020).

In the prior study characterizing the impact of the loss of function of *Sirt1* on metabolic homeostasis, the dysfunction observed was progressive in nature. While larvae and young adults are immediately obese, hyperglycemia and insulin resistance set in and become worse with increasing age (Reis *et al.* 2010; Palu and Thummel 2016). To ensure an appropriate set of conditions with respect to diet and age, we performed a preliminary analysis on 37 DGRP strains at one or two weeks of age, and under fasted or ad libitum fed conditions (Table S1). These time points correspond to conditions in the original study where differences in the metabolic state of the flies corresponded to differences in glycemia (Palu and Thummel 2016). We wished to select a time point at which hyperglycemia was detectable, but also differed enough between strains to allow us to identify genetic variation associated with that heterogeneity.

Three samples were collected for glucose measurements in each strain and condition. Glucose levels vary across genetic background for each of the conditions being tested (Figure 1A-D). Average glucose for each strain is significantly correlated between one and two week fasted flies (R = 0.53, P = 4E-03), between one-week-old fed and fasted flies (R = 0.45, P = 0.020), and between one-week-old fed and two-week-old fasted flies (R = 0.62, P = 2E-04) (Figure S5A-C). This supports glucose concentration as a consistent quantitative measurement. Interestingly, evidence for correlation with two-week-old flies fed ad libitum is not as strong. While a significant correlation is still detected with one-week-old fasted flies (R = 0.43, P = 0.014), the correlation is not significant with two-week-old fasted flies (R = 0.25, P = 0.219) and one-week-old flies fed ad libitum (R = 0.34, P = 0.063) (Figure S5D-F). By two weeks of age, *Sirt1* loss-of-function flies are beginning to experience more severe symptoms of disease, and we expect to see variability in symptoms and behavior in response to those symptoms. Fed flies at two weeks may have more variable glucose because feeding behavior is a big contributor to glucose levels in flies that have not been subjected to a fast.

**Figure 1.**
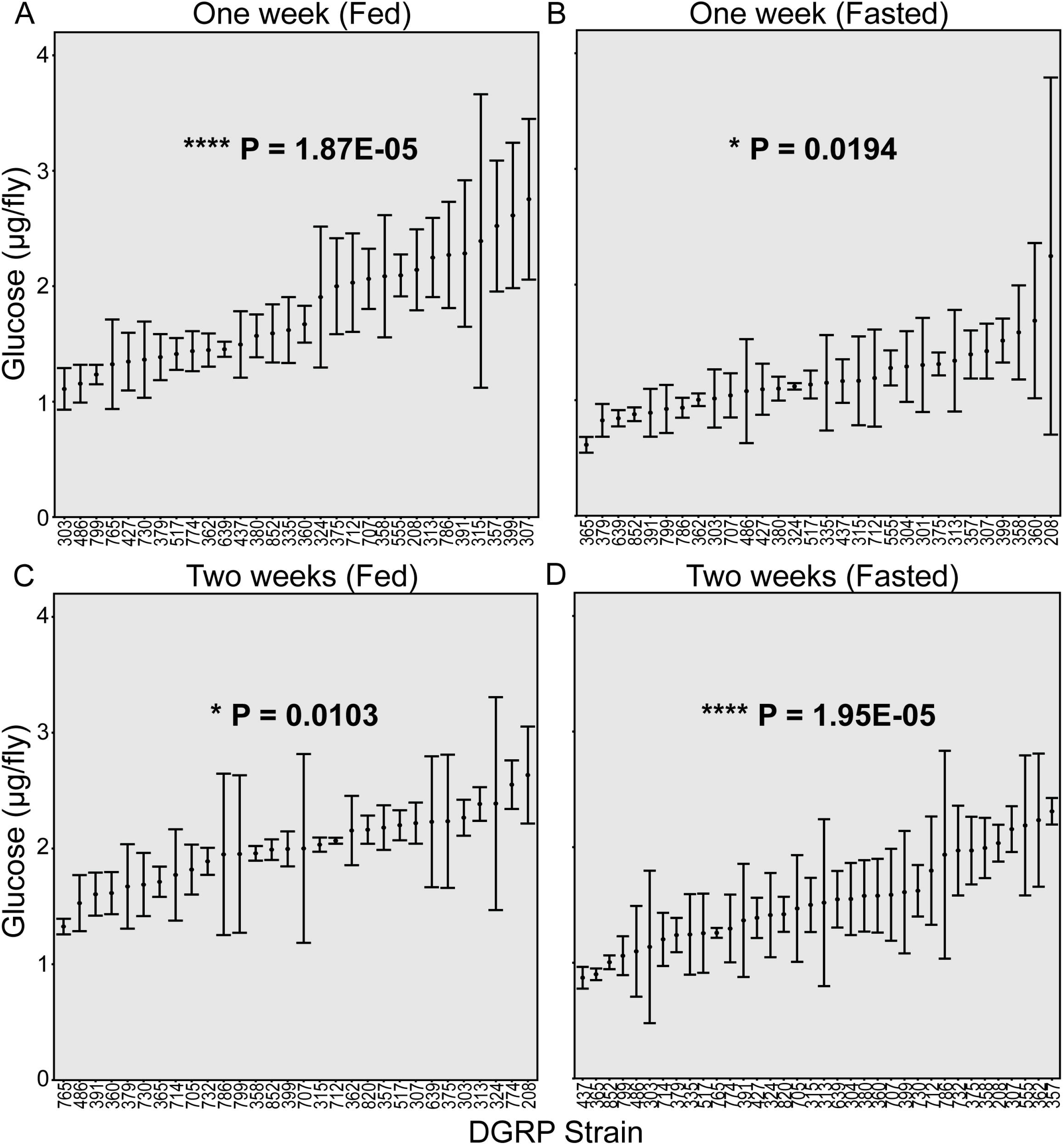
Glucose levels vary under a variety of environmental conditions. Glucose levels were measured in three samples for each of 30-36 strains under one of the indicated conditions: one week of adult age and fed ad libitum (N = 30 strains, P = 1.87E-05) (**A**), one week of adult age and fasted for 12-13 hours (N = 30 strains, P = 0.0194) (**B**), two weeks of adult age and fed ad libitum (N = 30 strains, P = 0.0103) (**C**), and two weeks of adult age and fasted for 12-13 hours (N = 36 strains, P = 1.95E-05) (**D**). Mean glucose concentrations are indicated, with error bars indicating standard deviation. DGRP strain or RAL numbers are indicated along the X-axis. P-values were calculated using one-way ANOVA incorporating all individual measurements comparing DGRP strain with glucose concentration. Adult flies were collected within 2-3 days after eclosion from the pupal case and aged to the indicated time points. * P < 0.05, **** P < 5E-05.

To identify the conditions under which the impact of genetic background was the strongest, we performed a one-way ANOVA test that included all data points collected. We found that while there is a significant association between glucose levels and genetic background under all conditions (p<0.05), this effect is most pronounced in the one-week-old flies fed ad libitum (P = 1.87E-5) and in the two-week-old fasted flies (P = 1.95E-5) (Figure 1A,D). Because fasting reduces possible intra-strain variation caused by food in the gut, two weeks fasted was selected for the full screen.

We examined protein concentrations in 90 samples from the first 30 strains collected at 2 weeks fasted to ensure that any variation we observe in glucose is not due to differences in body size. Protein levels do not significantly vary across the DGRP (P = 0.63, Figure S6A), nor do protein levels correlate with glucose levels in individual samples (R = 0.04, P = 0.6577, Figure S6B). We conclude that the variation we observe in fasting glucose levels are due to differences in circulating glucose, and not to differences in body size.

### Genome-wide association analysis identifies candidate modifiers of Sirt1i-associated hyperglycemia

Using the conditions determined in the preliminary screen, we proceeded to cross the donor strain with the remaining 149 DGRP strains (Figure 2, Tables S2,S3). We found a significant effect of genetic background on glycemia in the R4>Sirt1i flies (P < 2E-16) using one-way ANOVA including all data points for each strain (N = 555). Individual glucose measurements ranged from 0.306 μg/fly to 3.416 μg/fly (Table S2), while average concentrations ranged from 0.395 μg/fly (RAL 801) to 2.438 μg/fly (RAL 357) (Figure 2, Table S3).

**Figure 2.**
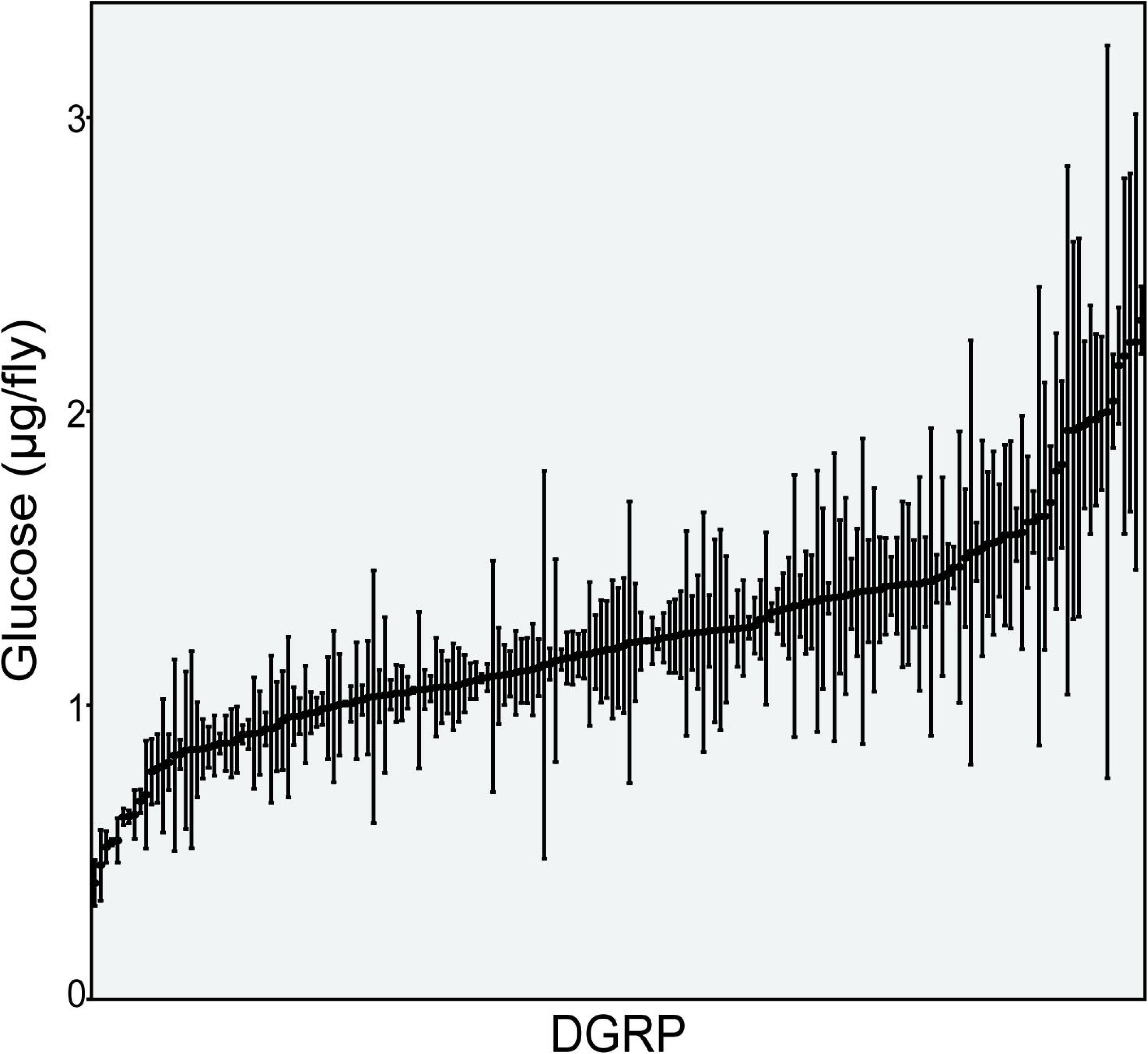
Glucose levels are significantly affected by genetic background. Glucose levels were measured in three samples for each of 185 strains at two weeks of age after a 12 hour fast. Adult flies were collected within 2-3 days after eclosion from the pupal case and aged an additional 9-11 days prior to fasting (11-14 days post-eclosion). Flies were collected after the overnight fast at 12-15 days post-eclosion. Mean glucose concentrations are indicated, with error bars indicating standard deviation. P-values were calculated using one-way ANOVA incorporating all individual measurements comparing DGRP strain with glucose concentration (P < 2E-16).

To identify genetic polymorphisms that may be responsible for this observed variation in glycemia, we performed a genome-wide association analysis. Average glucose level for each strain was used as a quantitative phenotype to test for association with polymorphisms in the DGRP. A total of 3,636,891 variants were tested for the *R4*>*Sirt1i* model across 186 lines. This analysis as a result is insufficiently powered for candidates to remain statistically significant after multiple testing corrections. Instead, the focus is on identification of candidate modifiers and pathways that can be validated through further study and that will provide the basis for future projects. This approach has been quite successful in previous studies (Chow *et al.* 2013, 2015, 2016; Palu and Chow 2018; Lavoy *et al.* 2018; Palu *et al.* 2019, 2020; Talsness *et al.* 2020).

Because the analyzed F1 hybrids in this case were male and inherited their × chromosome from the donor strain and not the DGRP strain, we do not include any X-linked variants in the resulting candidate modifiers. Using an arbitrary cut-off of P < 10^−4^, we identified 237 polymorphisms on the second and third chromosomes (Table S4). Of these 237, 62 were not considered further as they were not within +/− 1 kb of a candidate gene. The remaining 175 polymorphisms are associated with a total of 161 candidate genes (Table S5). 100 of these polymorphisms are intronic, 29 are exonic with 6 producing non-synonymous changes to the peptide and one a start-gain, 10 are located in the 5’ or 3’ untranslated regions, and 36 are within 1 kb up or downstream of the candidate gene (Table S4). Of note in this analysis is that the results were not filtered for allele frequency > 0.05. This was a concerted choice; several of the most interesting candidates, including *CG5888* and *ilp5*, would otherwise have been left out of the analysis. The number of total variants analyzed drops from 237 to 90, and the number of candidate genes associated at P < 10^−4^ drops from 161 to 75. While this, along with the use of a low stringency p-value cut-off, increases the probability of false positives, it likewise increases our power when performing pathway enrichment. Validation of candidate genes with low minor allele frequencies in later studies will distinguish true positives from false positives.

One concern with using an RNAi model to reduce *Sirt1* expression is that the modifiers identified might be specific RNAi efficacy, rather than hyperglycemia. The modifiers could be altering the degree and efficiency of *Sirt1* knockdown, so that hyperglycemia is actually correlating with the amount of *Sirt1* expression that is achieved. If this were the case, we would expect to see the top candidate modifiers associated with RNAi machinery or the efficiency of the GAL4/UAS system. This does not appear to be the case, either from a single-gene function perspective or when looking at enriched gene categories (Tables S5 and S6). Furthermore, the GAL4/UAS system is commonly used to model disease in the DGRP, and the candidates identified have always been unique to the disease and, at times, even the specific model in question (He *et al.* 2014; Chow *et al.* 2016; Lavoy *et al.* 2018; Palu *et al.* 2019; Talsness *et al.* 2020). All of this suggests that the candidate genes identified through this screen are modifying *Sirt1*-associated hyperglycemia directly rather than altering the degree of *Sirt1* knockdown.

### Candidate modifiers of Sirt1 are involved in basic metabolic processes, the immune response, and the regulation of neuronal communication

Because loss of *Sirt1* in the fat body alters glucose metabolism and insulin sensitivity in the organism, we expected modifiers of hyperglycemia to impact pathways linked to central carbon metabolism as well as external processes that influence secretion and signaling of hormones such as insulin. To determine if this is the case, we examined the individual functions of the top GWA candidates and looked for pathways and processes that are enriched in this list. While we attempted first to do this through Gene Ontology analysis of our top candidates, we found no significantly enriched terms. We therefore utilized individual known physical interactions and GO term enrichment through GSEA to highlight likely candidate pathways.

#### Analysis of Candidate Modifiers

We expected our top candidates to include genes that function in pathways or processes related to *Sirt1* regulation or activity. Among the most interesting candidates are those involved in NAD metabolism. We identified two genes whose products are part of the NADH dehydrogenase component of Complex I in the electron transport chain (*NDUFS3* and *ND-PDSW*) (FlyBase Curators 2008; Gaudet *et al.* 2011). There are also two NADP kinases, enzymes involved in the generation of NADP from NAD (*CG33156* and *CG6145*) (Gaudet *et al.* 2011). *DUOX*, an NADPH oxidase, passes electrons from NADPH to oxygen, generating hydrogen peroxide and altering the redox balance of the cell (Gaudet *et al.* 2011; Anh *et al.* 2011). As Sirt1 utilizes NAD as a cofactor during its enzymatic reaction, altering the balance of NAD in the cell through differential regulation of these enzymes could further impact the activity of other pathways that require NAD as an electron carrier, or exacerbate the phenotypes associated with *Sirt1* loss-of function (Nogueiras *et al.* 2012). We also identified two genes that have previously been implicated in the regulation and/or extension of lifespan: *sugb* and *CG42663* (Landis *et al.* 2003; Paik *et al.* 2012). While the role of *Sirt1* in lifespan extension is still contested, the identification of other genes implicated in this process suggests shared functions or pathways.

Genes involved in central glucose metabolism as well as insulin signaling are also candidate modifiers. MFS5 acts as a transporter of both glucose and trehalose for the uptake of these sugars from circulation (McMullen *et al.* 2021). Glucose-6-phosphatase (*G6P*) is the last rate-limiting step in both gluconeogenesis and glycogenolysis, which are used to generate glucose for release into the body during fasting (Gaudet *et al.* 2011; Lizák *et al.* 2019). These two genes directly regulate circulating glucose levels. Candidates involved in other metabolic pathways include *CTPSyn*, which encodes the rate limiting step in cytidine synthesis, the very long chain fatty acid ligase *bgm*, the mannosidase *Edem2*, and the oxoglutarate dehydrogenase complex subunit *CG33791* (Kang and Ryoo 2009; Gaudet *et al.* 2011; Jang *et al.* 2015; Sivachenko *et al.* 2016; Zhou *et al.* 2019). Partially responsible for regulating general metabolic flux through these various pathways is insulin. Interestingly, a top candidate is *ilp5*, one of several insulin-like peptides expressed in the insulin-producing cells (IPCs) in the *Drosophila* brain (Géminard *et al.* 2009). Our analysis also identified *IA-2*, a phosphatase involved in ilp secretion, *CG4168*, an uncharacterized gene whose closest human orthologue *IGFALS* encodes a protein that binds to and stabilizes IGF proteins in circulation, and *wrd*, a subunit in the protein phosphatase PP2A that negatively regulates insulin and TOR signaling (Boisclair *et al.* 1996; Kim *et al.* 2008; Hahn *et al.* 2010).

Other potential modifiers of insulin stability and signaling in circulation are *dally*, *cow*, and *Hs3st-A*. Both *dally* and *cow* encode heparin sulfate proteoglycans, while *Hs3st-A* encodes an O-sulfotransferase that acts on these proteoglycans (Filmus and Selleck 2001; Gaudet *et al.* 2011; Chang and Sun 2014). Previous work has demonstrated an impact of the enzyme heparanase, which cleaves heparan sulfate, on diabetic autoimmunity and complications such as nephropathy (Rabelink *et al.* 2017). While this is more peripheral to the central insulin signaling pathway in *Drosophila*, it highlights the utility of such factors in altering disease processes in subtle ways.

Another interesting group of candidates are those associated with neuronal development and function. Several members of the defective proboscis extension response (*dpr*) family were represented in our list (*dpr2*, *dpr6*, and *dpr13*) along with the dpr-interacting protein *DIP*-*eta*. The *dpr* gene family is collectively associated with synapse organization and function, as are the candidate genes *fife*, *CG32373*, and *atilla* (FlyBase Curators *et al.* 2004; Kurusu *et al.* 2008; Carrillo *et al.* 2015; Bruckner *et al.* 2017). We also noted candidates involved in neuropeptide signaling (*rk* and *RYa-R*), voltage-gated potassium channels and their regulation (*CG5888* and *CG1688*), and axon guidance (*tutl*, *CadN2*, *CG34353*, and *sbb*) (Rao *et al.* 2000; Luo *et al.* 2005; Prakash *et al.* 2005; Al-Anzi and Wyman 2009; Ida *et al.* 2011; Gaudet *et al.* 2011). The IPCs in *Drosophila* are actually neuroendocrine cells located in the brain, as are the AKH-producing cells responsible for secreting the glucagon-like hormone AKH (Géminard *et al.* 2006). The secretion of insulin is therefore dependent upon the correct development, connection, and signaling of neuronal cells.

The immune response is another generally enriched category of modifier genes. Several members of the nimrod family of immunoglobulins (*NimB2*, *NimC1*, and *NimC3*) were identified by GWA. All are implicated in the innate immune response, with *NimC1* and *NimC3* in particular having roles in phagocytosis (Somogyi *et al.* 2010). In response to insecticides, *LRR* regulates the immune response through NF-kappaB, whose activation is an early protective event in the progression and pathology of diabetes (Prisco *et al.* 2013; Irvin *et al.* 2018). Two lysozyme enzymes with links to bacterial defense (*LysX* and *CG7798*) highlight the role of oxidative stress and redox homeostasis in the innate immune response (FlyBase Curators 2008). As Sirt1 has roles in regulating the response to oxidative stress, we looked for other genes with similar functions (Brunet *et al.* 2004). As described above, *DUOX* plays a part in regulating redox homeostasis through the production of hydrogen peroxide (Anh *et al.* 2011). *CG42331* encodes a peroxidase that appears to be strongly enriched in the pupal fat body, and *cyp28a5* encodes an oxidoreductase that, similar to LRR, is involved in the response to insecticides (FlyBase Curators *et al.* 2004; Graveley *et al.* 2011; Gaudet *et al.* 2011). It is now believed that Type I and Type II diabetics both suffer at least to some degree from autoimmunity (Candia *et al.* 2019). Exploring the direct and indirect connections of *Sirt1* to the immune response and oxidative stress directly is an interesting avenue for future direction.

#### Physical Interaction Network

We generated a network of physical interactions among the 161 candidate genes identified above. These were identified and visualized using Cytoscape software with the GeneMania plugin (Shannon *et al.* 2003; Montojo *et al.* 2010). The products of 37/161 candidate genes were found to physically interact with at least one other candidate gene product with no more than one bridging node represented by a non-candidate gene (Figure 3A). This high degree of interaction suggests that the modifiers identified in this screen are indeed functioning through shared processes.

**Figure 3.**
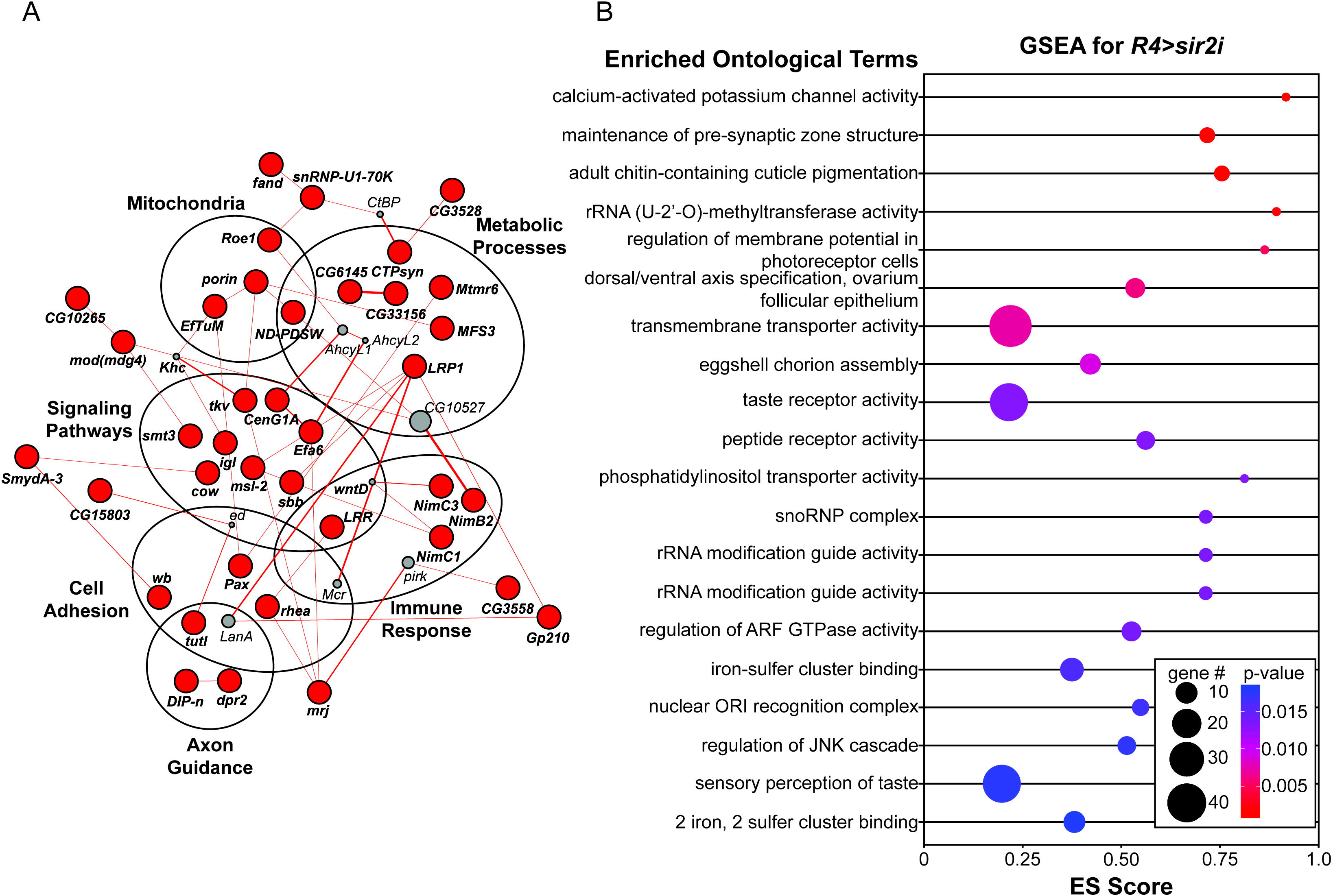
Immune responses, neuronal function, and basic metabolic processes are overrepresented in GWA candidate modifiers of hyperglycemia. (**A**) *R4*>*Sirt1i* modifier network, as plotted by the GeneMANIA plugin in Cytoscape (Shannon *et al.* 2003; Montojo *et al.* 2010). Significant candidate modifiers are indicated in red, with physical interactions indicated by connecting red lines. Thicker lines indicate stronger evidence for the interaction. Encircled genes share common pathways or functions. Interacting genes outside of the candidate modifier list are indicated in gray. (**B**) Top 20 significant ontological categories as identified by GSEA. Categories are arranged from most significant on top to least significant along the y-axis. P-values are indicated by red-to-blue gradient, with red the lowest p-values and blue the highest P-values. Enrichment score (ES) for each category is plotted along the x-axis. Gene number identified in each category is indicated by dot size.

Focusing then on the 37 genes involved in physical interactions, we identified several broad functional categories that could influence glucose homeostasis in the fly. The most obvious category are enzymes that catalyze steps in basic metabolic pathways (N = 7). This includes the NAD kinases (*CG6145* and *CG33156*) and one NADH dehydrogenase (*ND*-*PDSW*) discussed above (Gaudet *et al.* 2011). Other metabolic candidate modifiers include *MFS3* and *CTPsyn*, the rate limiting enzyme in the production of the nucleotide Cytidine triphosphate (Zhou *et al.* 2019). Both of these enzymes function in pathways critically dependent upon or feeding into central carbon metabolism, and their inclusion as candidate modifiers of hyperglycemia supports a role for those secondary metabolic pathways as a sink for increased circulating glucose. *Mtmr6* encodes a phosphatidylinositol phosphatase, a key enzyme in several signaling pathways, including insulin signaling (Gaudet *et al.* 2011). Also interesting is the gene *LRP1*, which is orthologous to human LDL receptor related protein 1. In additional to its role in lipid homeostasis, *LRP1* has also been implicated in Alzheimer’s disease, for which metabolic disease and obesity are risk factors (Kang *et al.* 2000; Anstey *et al.* 2011).

Curiously, several of the genes highlighted in this analysis also happen to localize specifically to the mitochondria (N = 4). *Roe1* and *porin* are both transporters involved in the import of molecules into the mitochondria (Komarov *et al.* 2004; Jana Alonso *et al.* 2005; FlyBase Curators 2008; Gaudet *et al.* 2011). The NADH dehydrogenases *ND*-*PDSW* and *NDUFS3* both function in the mitochondria as well as part of Complex I in the electron transport chain (Jana Alonso *et al.* 2005). Closer examination of the top GWA candidates reveals additional mitochondrial localization candidates including the amino acyl tRNA synthetase *GlyRS*, the membrane bound regulator of protein kinase A (*pkaap*) and the translation elongation factor *mEFTu1* (Gaudet *et al.* 2011; Lu *et al.* 2016). In adult metabolic homeostasis, central carbon metabolism is generally used to fuel the electron transport chain in the mitochondria and generate ATP for the cell (Barry and Thummel 2016). Altering the activity of this essential downstream pathway could have a clear impact on glucose utilization and disease progression in diabetes.

Similar to our examination of top candidates, our physical interaction map highlighted the immune response (*NimC1*, *NimC3*, *NimB2*, and *LRR*) and neuronal function. *Tutl*, *dpr2*, and *DIP*-*eta*, and *wb* are all involved in synapse organization and axon guidance, while *Pax* and *rhea* are involved in focal adhesion (Delon and Brown 2009). The identification of genes important for cellular communication suggests that some of the modifiers identified in this study have roles in tissues other than the fat body, such as the IPC and APC neurons. This is an important avenue of future exploration.

#### Gene Set Enrichment Analysis (GSEA)

In the second approach, we performed GSEA analysis to identify gene ontology terms for which associated variants are enriched. Unlike traditional GO analysis, which relies upon a set of genes based on a P-value cutoff, GSEA examines the entire gene set (Dyer *et al.* 2008). For each defined GO category, GSEA determines whether the members of that category are randomly distributed throughout the ranked gene list provided or if they are enriched for the lower p-values found at the top of that list. GO categories enriched at the top of the list describe important functions of the gene set. GSEA identified 52 significantly associated gene sets (≥ 3 genes) with positive enrichment scores at a p-value of <0.05 (Table S6, Figure 3B). The top two gene sets implicate neuronal function and communication in the *Sirt1i*-associated hyperglycemia phenotype: calcium-activated potassium channel activity (GO:0015269, P = 1.1E-3) and maintenance of presynaptic active zone structure (GO:0048790, P = 1.2E-3). Similar categories can be found through the list of significantly associated gene sets, including dendrite morphogenesis (GO:0048813, P = 0.049), which represents the largest group of genes at N = 119 and contains two of the top GWA candidates (*slit* and *fruitless*). Coupled with the neuronal genes identified in our physical interaction network, this suggests that function in the neuroendocrine cells could play a big role in glucose homeostasis in the fat body.

Also enriched are taste receptor activity (GO:0008527, P = 0.013) and sensory reception of taste (GO:0050909, P = 0.018). These categories highlight a yet unconsidered factor that must be taken into account with metabolic phenotypes: that of feeding and diet. While all flies were collected under identical conditions and were maintained on the same diet, it is nonetheless possible that some may simply be eating more due to differences in the sensing of satiation or to differences in perception of taste. These differences in perception and consumption can have detectable impacts on metabolic phenotypes (May *et al.* 2019).

As expected, we also see evidence of general metabolic processes. Some, like alpha,alpha-trehalase activity (GO:0004555, P = 0.043) and phosphatidylinositol transporter activity (GO:0008526, P = 0.013) have direct links to glucose metabolism and insulin signaling. Others, such as oxysterol binding (GO:0008142, P = 0.026), glutamate biosynthetic process (GO:0006537, P = 0.043), and isoprenoid biosynthetic process (GO:0008299, P = 0.048) function more peripherally to carbon metabolism and are likely influencing hyperglycemia by their general contribution to physiological homeostasis.

Another interesting group of GO categories highlighted by GSEA are RNA processing functions. rRNA (uridine-2′-O-)-methyltransferase activity (GO:0008650, P = 2.4E-3) is the fourth most associated category as ranked by p-value, and others such as snoRNA binding (GO:0030515, P = 0.014) reiterating this function. The presence of RNA processing categories is of particular interest because three of the top candidate genes by GWA are splicing factors (*bru1*, *fand*, and *snRNP*-*U1*-*70k*) (Park *et al.* 2004; Oas *et al.* 2014; Spletter *et al.* 2015). While it is unclear how rRNA or mRNA processing may directly or indirectly influence glucose homeostasis in particular, the identification of this process through several different methods of analysis is striking and worth further exploration.

### Functional analysis of candidate modifier genes

To confirm the roles of our candidate genes in regulating glucose homeostasis, we elected to test the impact of loss of modifier expression for 16 of the most significant candidates for which we were able to obtain transgenic RNAi lines (Table 1). We crossed the RNAi strains targeting each of these modifiers into the *R4*>*Sirt1i* line, aged the resulting progeny for 2-3 weeks, and measured glucose in fasted males. We also measured protein levels as a control. Knockdown of modifier genes did not significantly alter protein levels as compared to a genetically matched control (Figure S7). Knockdown of the genes *CG4168*, *CG5888*, and *uif* resulted in suppression of the hyperglycemia phenotype, with a significant decrease in glucose content per fly compared to controls expressing *R4*>*Sirt1i* (Figure 4). These results demonstrate that many of the top GWA candidate modifiers are capable of modifying the hyperglycemia phenotypes associated with the *R4*>*Sirt1i* model of diabetes.

**Figure 4.**
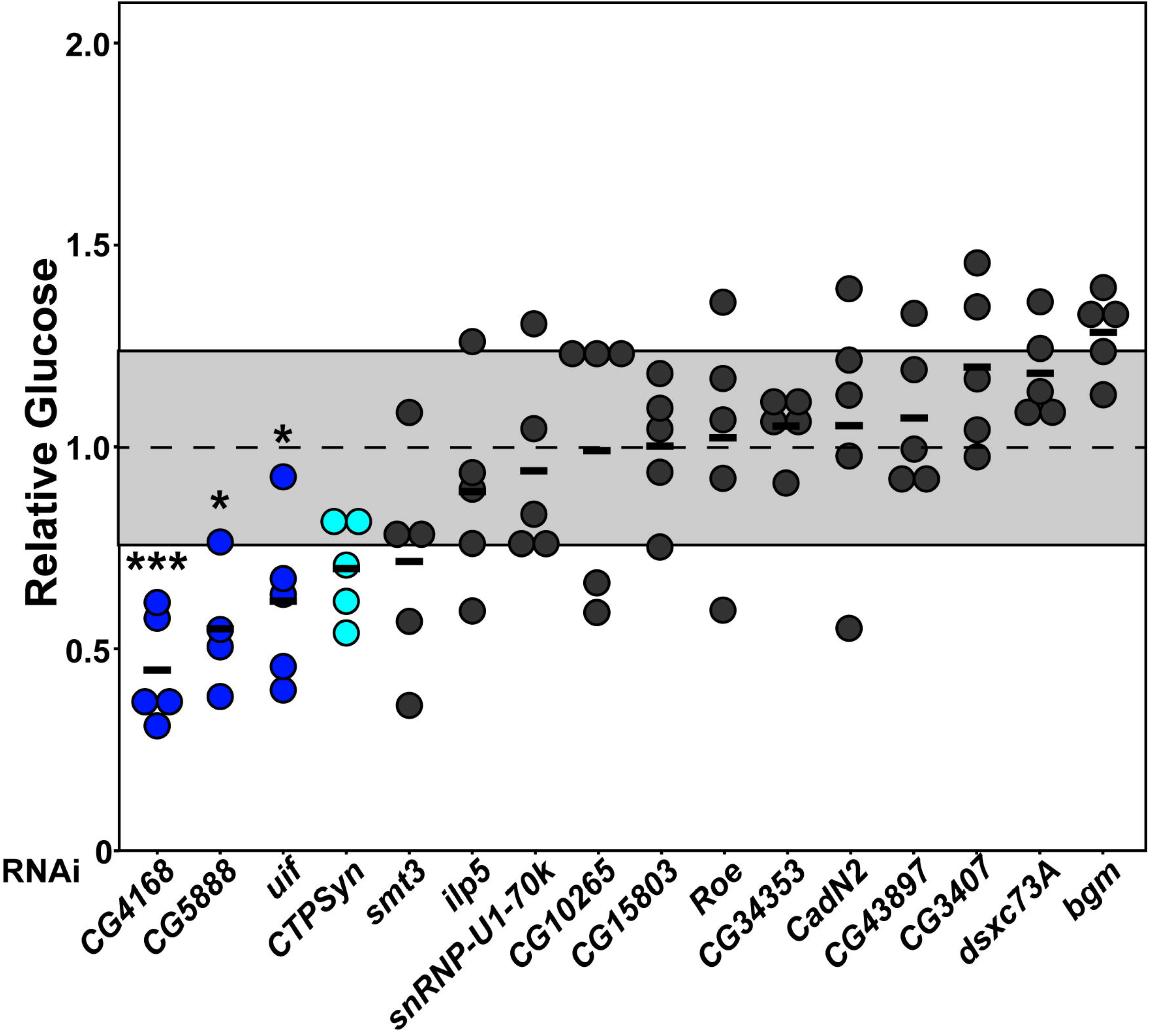
Loss of candidate gene expression suppresses hyperglycemia in the R4>Sirt1i model. RNAi against candidate modifiers was expressed under the control of *R4*-*GAL4* in the *R4*>*Sirt1i* model. Glucose level for each sample was normalized to the levels in a genetically matched control line crossed into the *R4-Sirt1* line. Average *R4*>*Sirt1i* control glucose levels after normalization are indicated by a dotted line at 1.0, with standard deviation highlighted by the gray box. Whole fly glucose concentration was quantified for N = 4-5 samples per strain, each consisting of 5 flies and individually plotted along the y-axis. Knockdown of *CG4168, CG5888*, or *uif* significantly reduces glucose concentrations in the *R4*>*Sirt1i* model of hyperglycemia compared to controls (blue). Loss of *CTPSyn* does not significantly alter glucose levels, but a trending decrease in glucose levels were observed in several independent RNAi strains (light blue, data not shown). Loss of *smt3*, *ilp5*, *snRNP*-*U1*-*70k*, *CG10265*, *CG15803*, *Roe*, *CG34353*, *CadN2*, *CG43897*, *CG3407*, *dsxc73A*, or *bgm* do not produce a significant effect (dark gray). P-values were calculated using one-way ANOVA followed by Dunnett’s multiple testing correction. * P < 0.05, *** P < 0.001.

## DISCUSSION

Identifying and characterizing the genetic factors influencing the severity of diabetes is critical to early diagnosis. Prevention is still the best strategy available, and providing patients at high risk for complications with knowledge of that risk could prevent the worst symptoms from manifesting. It could even enable intervention before the progression of disease is irreversible.

In this study, we identified and analyzed a number of candidate modifiers of hyperglycemia in a previously characterized model of diabetes, *Sirt1* loss-of-function. We used the DGRP as an unbiased source of natural genetic variation for this screen. This is the first time a genetic model of metabolic dysfunction has been put to this use, as previous screens have either focused on dietary stress as a source of metabolic disease or on the impact of genotype on metabolic parameters under non-stressed conditions (Mackay *et al.* 2012; Ivanov *et al.* 2015; Nelson *et al.* 2016; Jehrke *et al.* 2018; Everman *et al.* 2019). We observed very little overlap in modifier candidates between our observations and these studies. This is consistent with previous work demonstrating that even when the observed phenotypes are similar or nearly identical, the overlap in modifiers is often small in different models of disease (Palu *et al.* 2019). One exception is a screen for the response to starvation resistance performed by Everman *et al.* in 2018. They identified *CG15803*, a transporter of unknown function that was also identified in our studies (Everman and Morgan 2018). While our preliminary analysis suggests that this gene does not function in the fat body, it is possible that it could have a role in another physiologically relevant tissue such as the IPCs or APCs. Indeed, *CG15803* appears to be most highly expressed in the head and CNS (Graveley *et al.* 2011). This is an interesting avenue of future exploration.

We did observe overlap in general gene categories with previous studies, even when direct overlaps were few. This was found to be true for modifiers of neuronal function. In a broad exploration of genetic variation in the nutrient response that looked at triglycerides, starvation resistance, mass, and glucose, *NimB3* was identified as a candidate modifier (Nelson *et al.* 2016). While *NimB3* was not identified in this screen, we did find *NimB2*, *NimC1*, and *NimC3*. *Fife*, which also has roles in synapse organization, was identified in this study and one for starvation resistance (Mackay *et al.* 2012). As mentioned above, both the IPCs and APCs are neuroendocrine cells found in or near the brain (Géminard *et al.* 2006). Maintenance of neuronal function would be critical to hormonal balance as a result. Furthermore, it is broadly acknowledged that metabolic homeostasis is also dependent upon feeding rate, over which the central nervous system has some sway (Owusu-Ansah and Perrimon 2014). The identification of neuronal genes in each analysis suggests that regulation of particular neuronal pathways and cells is critical to the maintenance of physiological homeostasis.

Given the prevalence of neuronal and sensory perception genes in the analysis, a concern could be raised for the role feeding rate could be playing in the variation of glucose across the DGRP. To assess this, we compared glucose with male feeding rate in a previous analysis (Table S3) (Garlapow *et al.* 2015). We saw no correlation whatsoever, indicating that while nutrient sensing may play a role in the response to Sirt1i-induced hyperglycemia, it is not the driving factor in the variation observed in this screen (Figure S8).

Another interesting category highlighted through several analysis methods is the innate immune response. While it has long been known that Type I diabetes is an autoimmune disorder, it has recently been acknowledged that Type II diabetics also display symptoms of autoimmunity (Candia *et al.* 2019). Furthermore, insulin resistance has frequently been associated with inflammation, and the presence of macrophages in the adipose tissue is a hallmark of obesity and diabetes (Wu and Ballantyne 2020). Modifiers associated with innate immunity serve therefore as validation to the study as a whole, and examination of these genes and their function in the context of the *Sirt1* loss-of-function model will be an intriguing focus of future research.

An important component of this study is the validation of top candidate modifiers using RNAi-mediated knockdown of gene expression. We obtained strains expression transgenic RNAi constructs targeting 16 of the most significant candidates (Table 1). We found that reduced expression of three candidates specifically in the fat body resulted in significant suppression of hyperglycemia: *CG4168*, *CG5888*, and *uif* (Figure 4). We also noted a consistent, though not significant, decrease in glucose for two independent RNAi constructs targeting *CTPSyn*, suggesting that this gene warrants further study (Figure 4, data not shown). The remainder of the genes had no significant or consistent impact on hyperglycemia in the model of *Sirt1* loss-of-function (Figure 4). There were no enhancing modifiers: loss of modifier expression did not lead to increased glucose levels for any of the tested genes. This could be because hyperglycemia is already quite strong in the *Sirt1* loss-of-function model, or because we simply did not hit on any enhancing modifiers. Of greater concern is the lack of any response for 13 of the 16 tested candidates. One explanation may be found in the large number of known neuronal genes identified in this analysis. This modifier RNAi screen specifically focused on the expression of the RNAi against the candidate genes in the fat body, where expression of *Sirt1* is also reduced. If, however, the function of a modifier gene is primarily concentrated in the IPCs, as with *ilp5*, reducing its expression in the fat body would have little effect on the disease phenotypes in question. We will examine the role of modifier genes not only in the fat body but in the IPCs, APCs, and other physiologically relevant tissues in future work.

Of immediate interest is of course the mechanism of action for the three suppressor genes that were confirmed by RNAi in the fat body. Of these, *CG5888* is also the top GWA candidate (P = 2.80E-07). While largely uncharacterized, *CG5888* has been identified as a component or activator of voltage-gated potassium channels by sequence homology (Gaudet *et al.* 2011). Intriguingly, it has also been implicated JNK signaling by a previous RNAi screen (Bond and Foley 2009). JNK signaling is commonly activated by cellular stress and can activate apoptosis. It also has important roles in the signaling pathways commonly used by the immune system (Bond and Foley 2009; Shlevkov and Morata 2012). A potential role for *CG5888* in immune pathways has yet to be explored and could be an exciting area of further discovery.

*Uninflatable* (*uif*) encodes a single pass transmembrane protein found on the apical membrane of epithelial cells and has been found to enable Notch signaling (Loubéry *et al.* 2014). It has also been found to exacerbate disease in a *Drosophila* model of muscular dystrophy (Kucherenko *et al.* 2011), and its closest human orthologue ELAPOR1 is a regulator of apoptosis and autophagy (Deng *et al.* 2010). Both of these processes are commonly disrupted through inappropriate activation in metabolic disease, and might provide some explanation for the impact of *uif* on *Sirt1i*-associated phenotypes (Bugliani *et al.* 2019).

Perhaps the most intriguing finding is *CG4168* as the modifier with the strongest impact on *Sirt1i*-associated hyperglycemia. The protein encoded by *CG4168* is of unknown function, but its closest human orthologue (*IGFALS*) encodes a serum protein that binds to insulin-like growth factors (IGF) in circulation (Boisclair *et al.* 1996). In mammals, association with IGFALS increases the half-life of insulin-like growth factors in the serum as well as their retention in circulation. While studies of IGFALS in mammals has not shown a role for it in regulating insulin signaling, the *Drosophila* ilp peptides are used for both IGF and insulin signaling activation (Géminard *et al.* 2006). It is possible that secretion of the factor encoded by *CG4168* from the fat body could increase ilp retention in circulation, whereas its loss could result in faster clearance of ilps from circulation. Under conditions that promote insulin resistance, such as the loss of *Sirt1*, it is possible that reduced ilp levels in the hemolymph could slow or prevent insulin resistance and hyperglycemia. Further exploration of the mechanisms behind the action of *CG4168* could reveal important insights into circulating insulin-binding factors and their role in diabetes.

In conclusion, we have identified a number of pathways and processes involved in the degree of hyperglycemia in a genetic model of diabetes. Examination of the candidate genes and pathways described above in this model as well as other models of metabolic dysfunction will shed new light on the mechanism by which insulin resistance and related complications disease onset, progression, and severity. Furthermore, the candidates identified as suppressors could serve as promising targets for therapeutics in diabetes and related metabolic disorders.

## DATA AVAILABILITY STATEMENT

Strains and stocks are available upon request, as is code for GSEA. Genomic sequence for the DGRP is available at http://dgrp.gnets.ncsu.edu/. Supplemental material is available at FigShare (https://figshare.com/s/2b0b0237de0f94139a4f).

## ACKNOWLEDGEMENTS

We thank Clement Y. Chow for contributions to the data analysis in this project, as well as reagents and comments. This research was supported by a Faculty Summer Research Grant through the Purdue University-Fort Wayne Office of Sponsored Projects to RASP as well as the Purdue University-Fort Wayne Department of Biology. KGO is supported by the NIGMS Genetics T32 Fellowship from the University of Utah (T32GM007464).

