## Supplemental Figures for "A natural genetic variation screen identifies insulin signaling, neuronal communication, and innate immunity as modifiers of hyperglycemia in the absence of *Sirt1*"

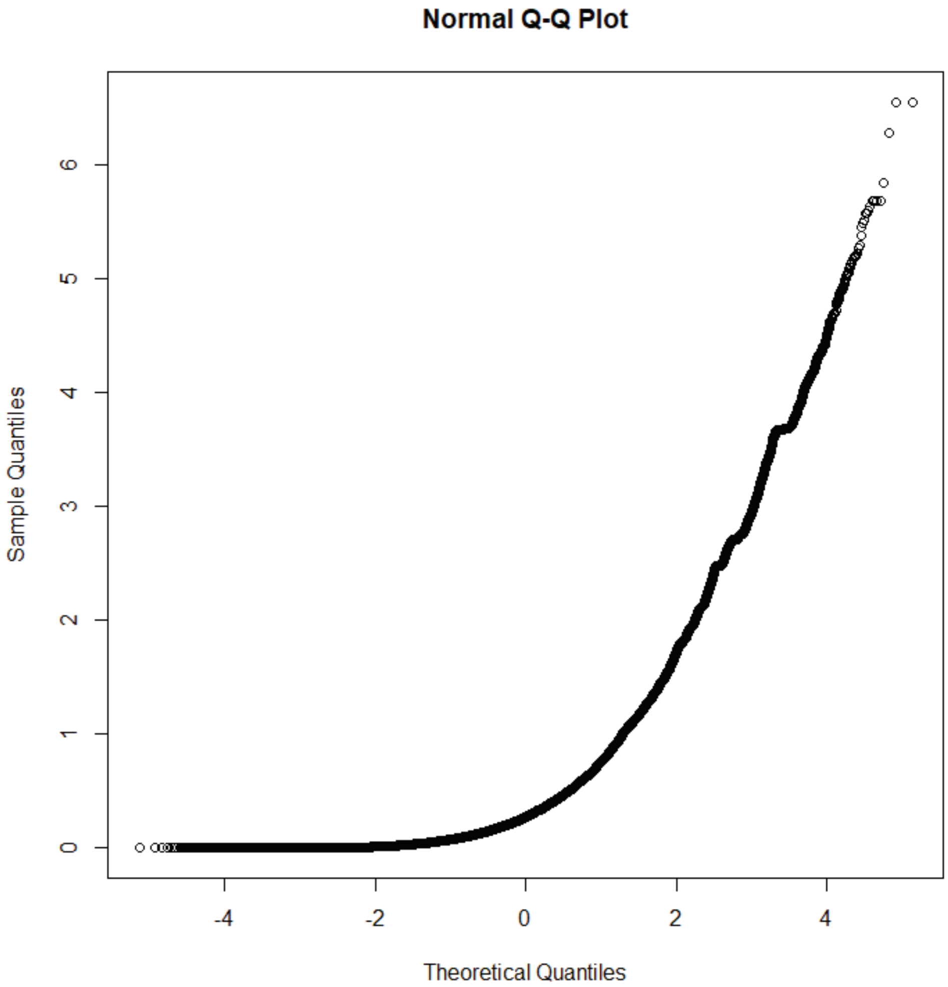


**Supplemental Figure 1. Distribution of datapoints from the Genome-Wide Association Analysis for *R4>sir2i* deviate from the LMM.** Quantile-quantile plots were generated for P-values (log10[P-value]) across the 4,202,640 variants tested for the R4>sir2i model of hyperglycemia. Calculated expected values are distributed along the x-axis, with observed values along with y-axis. The analysis is skewed on the low end of the plot, with more variants with very high, non-signficant p-values than would be expected by chance.

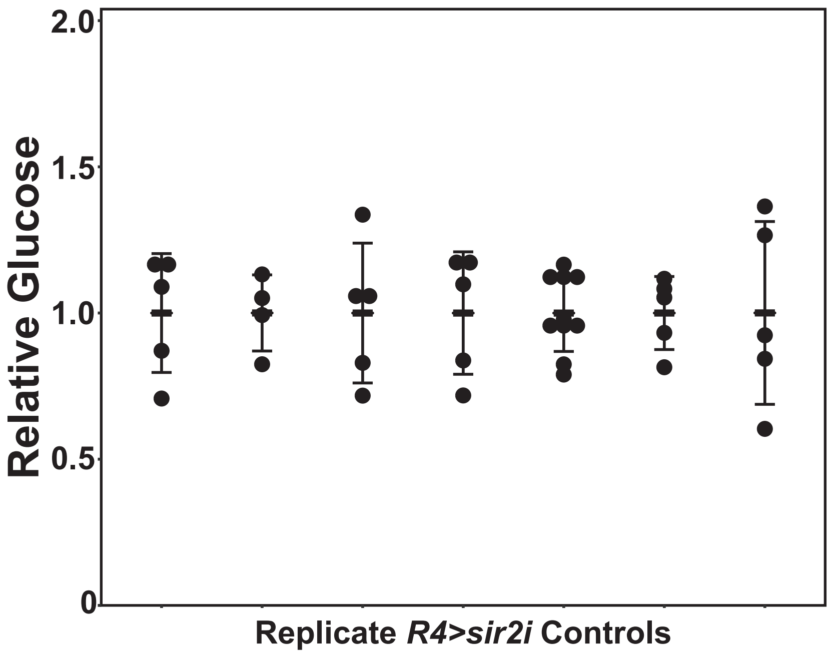


**Supplemental Figure 2. *R4>Sirt1i* controls do not vary in standard deviation from experiment to experiment.** Each replicate of the modifier RNAi experiment was compared to a genetically matched control expressing the R4>Sirt1i model of hyperglycemia. Each measurement is normalized to the average glucose levels for controls within the given replicate (relative average glucose level for all control strains = 1.0). Error bars represent standard deviation for each replicate control. Because results from several replicates were combined, the standard deviations from all controls were compared to ensure that the comparisons to RNAi strains were representative. There is no significant impact of replicate on standard deviation of glucose measurements (threshold $p<0.05$). All RNAi data in Figure 4 is compared to control replicate #3 in this figure.

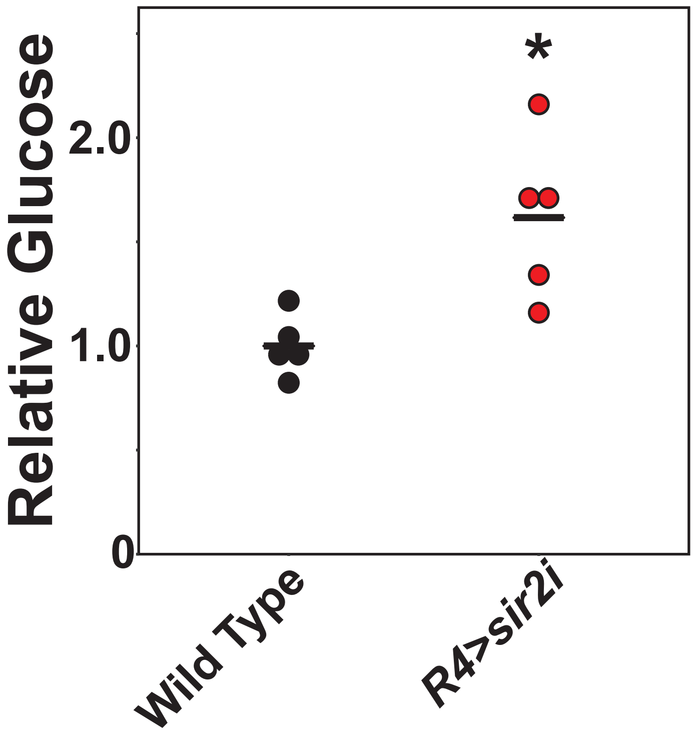


**Supplemental Figure 3. Loss of *sir2* in the fat body recapitulates hyperglycemia observed in *sir2* loss-of-function mutants.** The *R4-GAL4* driver is used to induce expression of an RNAi construct targeting *sir2* specifically in the fat body. A genetically matched control expressing only *R4-GAL4* serves as the wild-type strain. Male flies were aged two weeks and fasted overnight 16-20 hours prior to collection (N = 5 each genotype, with 5 flies in each sample). Glucose measurements are normalized to the average for the wild-type control. As observed in Palu and Thummel (2016), gluocse concentrations increase on average ~1.6 fold upon loss of *sir2* in the fat body as compared to genetically matched controls. * $P < 0.05$

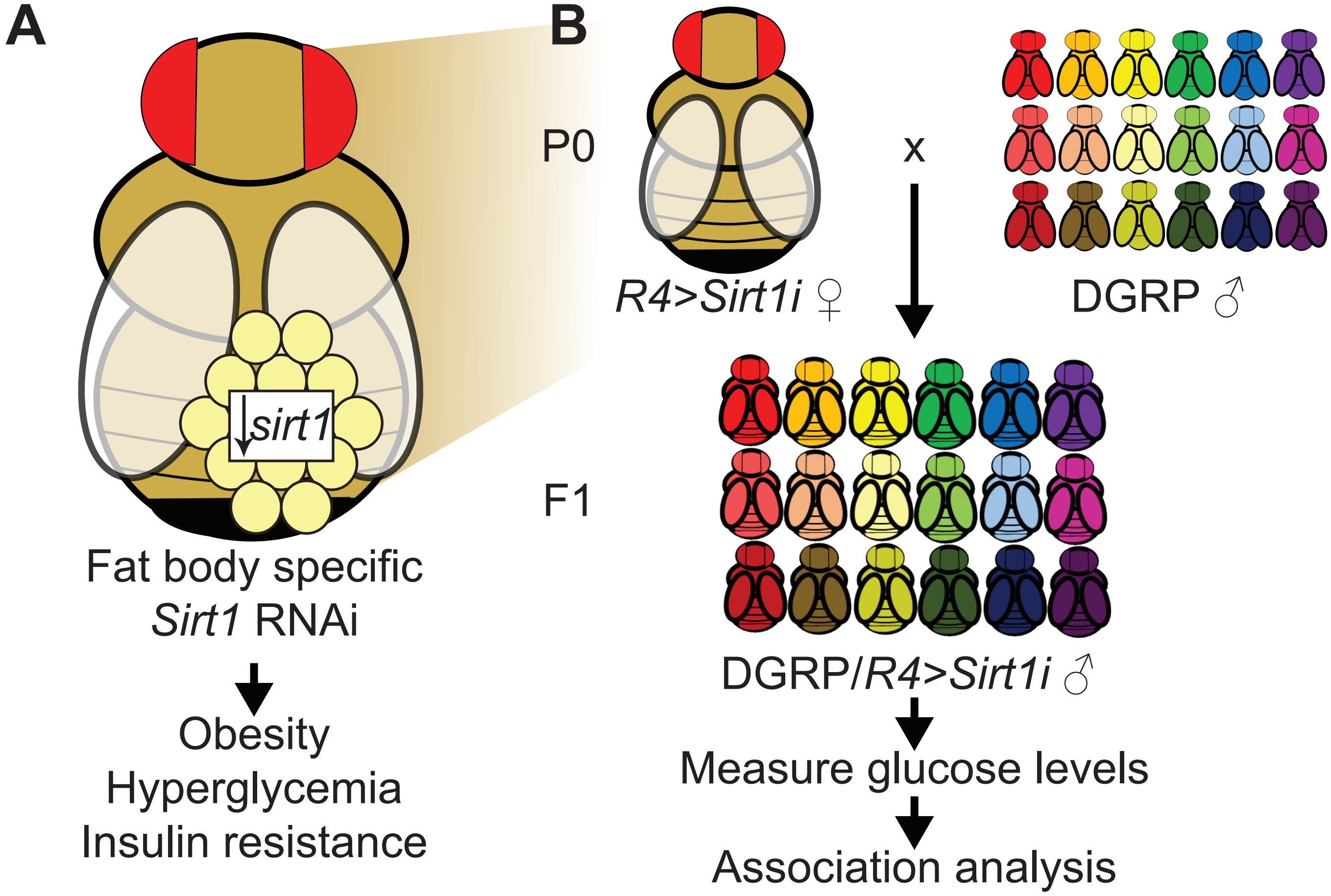


**Supplemental Figure 4. Model of Sirt1 loss-of-function.**
**(A)** *R4-GAL4* drives expression of *Sirt1* RNAi specifically in the fat body. This reduces *Sirt1* expression in that tissue and results in metabolic dysfunction (Palu and Thummel 2016). **(B)** F1 progeny of the *R4>Sirt1i* females and DGRP males will be assayed for glucose levels as a proxy for hyperglycemia.

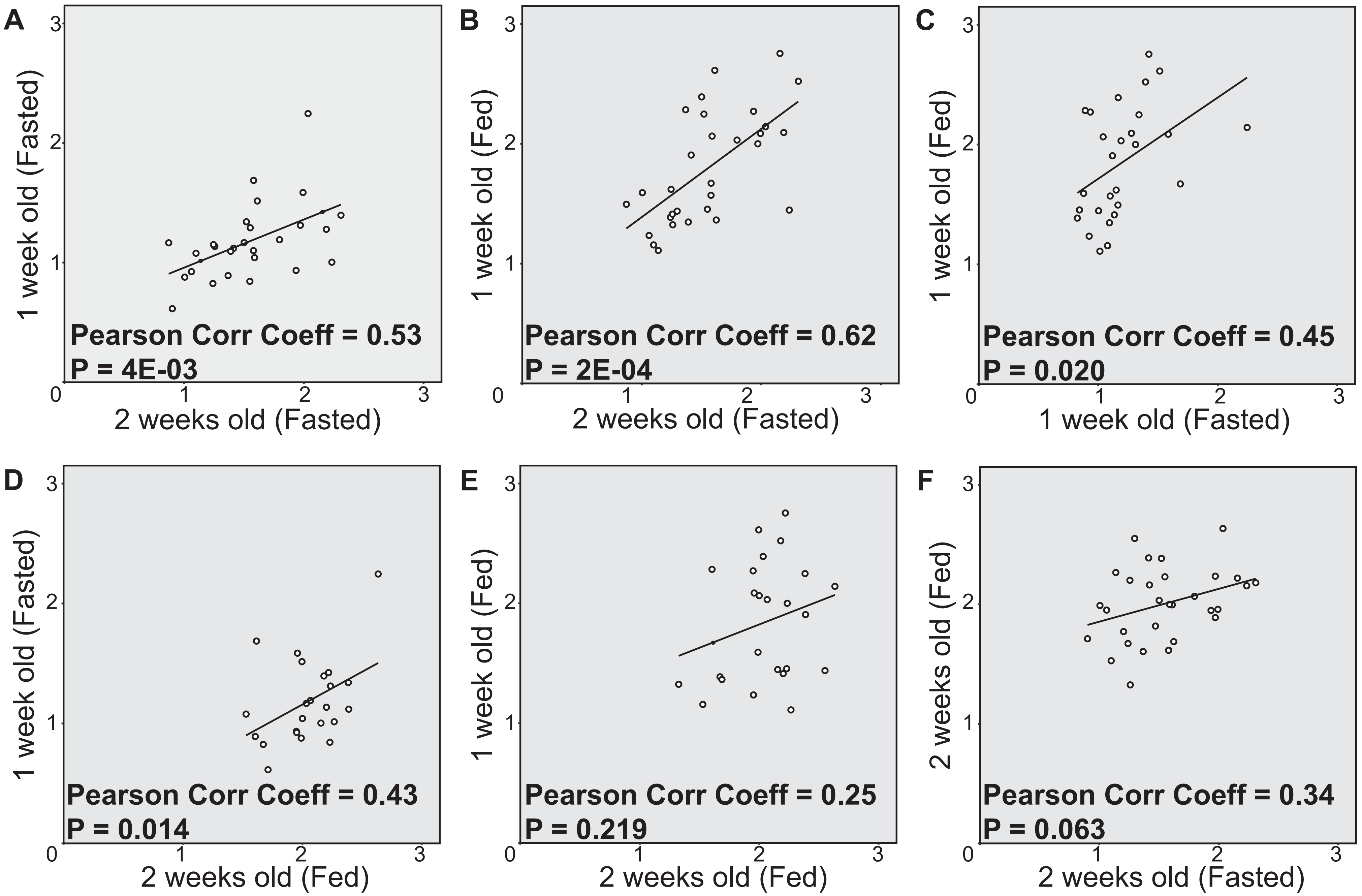


**Supplemental Figure 5. Glucose levels are correlated across different collection conditions.**
Correlation coefficients and p-values were calculated using a Pearson Correlation test. Glucose concentrations (μg/fly) across different DGRP strains were significantly positively correlated between two-week-old fasted flies and both one-week-old fasted flies **(A)** and one-week-old flies fed ad libitum **(B)**, as well as between both one-week-old fasted and one-week-old flies fed ad libitum **(C)** and two-week-old flies fed adlibitum **(D)**. The correlation was weaker and not statistically significant between two-week-old flies fed ad libitum and both two-week-old fasted flies **(E)** and one-week-old flies fed ad libitum **(F)**.

Two weeks (Fasted)

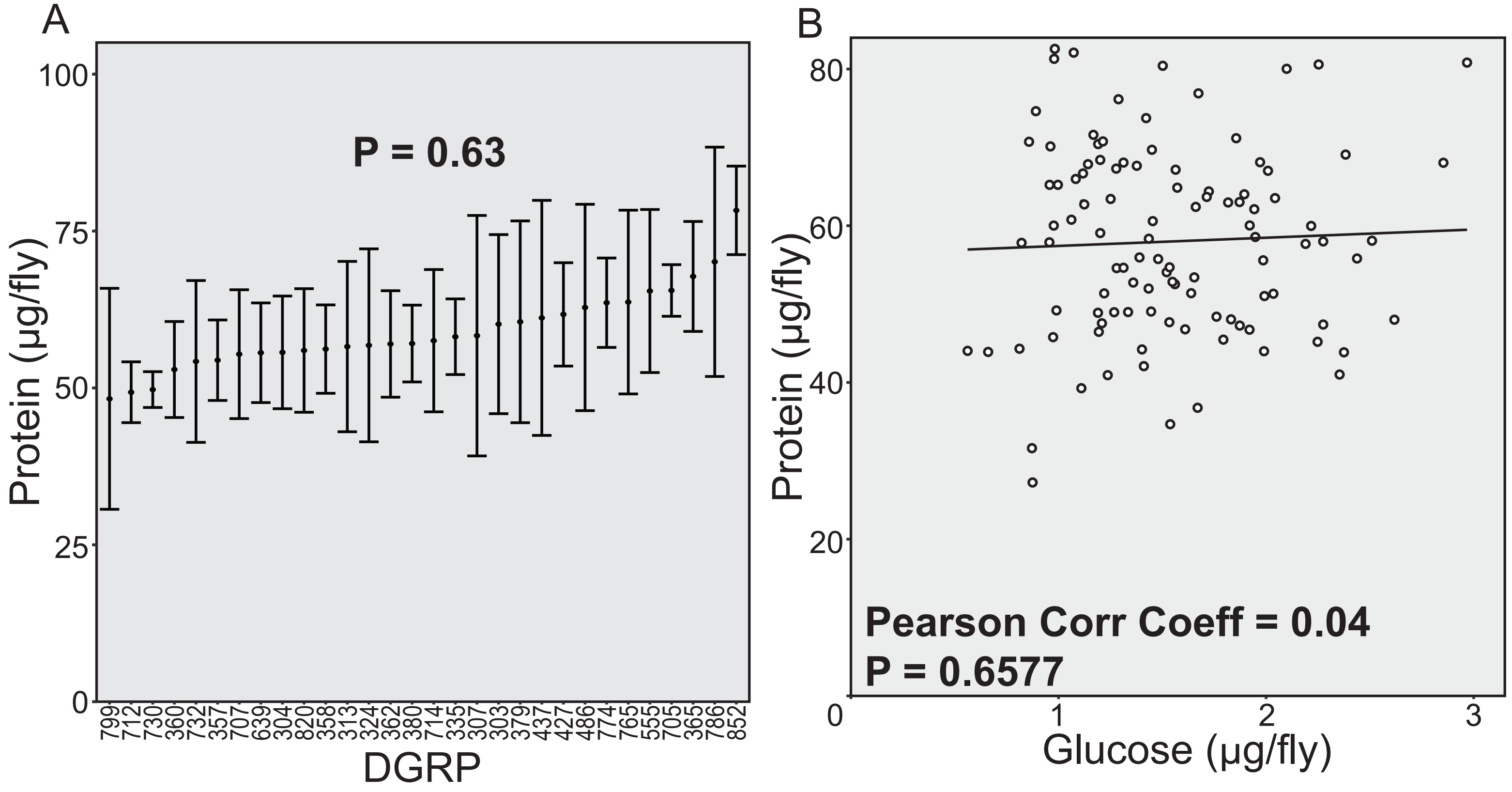


**Supplemental Figure 6. Protein levels do not impact variation in glucose levels.**
Protein and glucose were measured from the same isolates. Mean protein concentrations are indicated, with error bars indicating standard deviation. The P-value was calculated using one-way ANOVA incorporating all individual measurements. Protein levels were not significantly affected by genetic background **(A).** Protein concentration of individual samples was not correlated with glucose concentration of the same sample as determined by Pearson correlation test **(B)**.

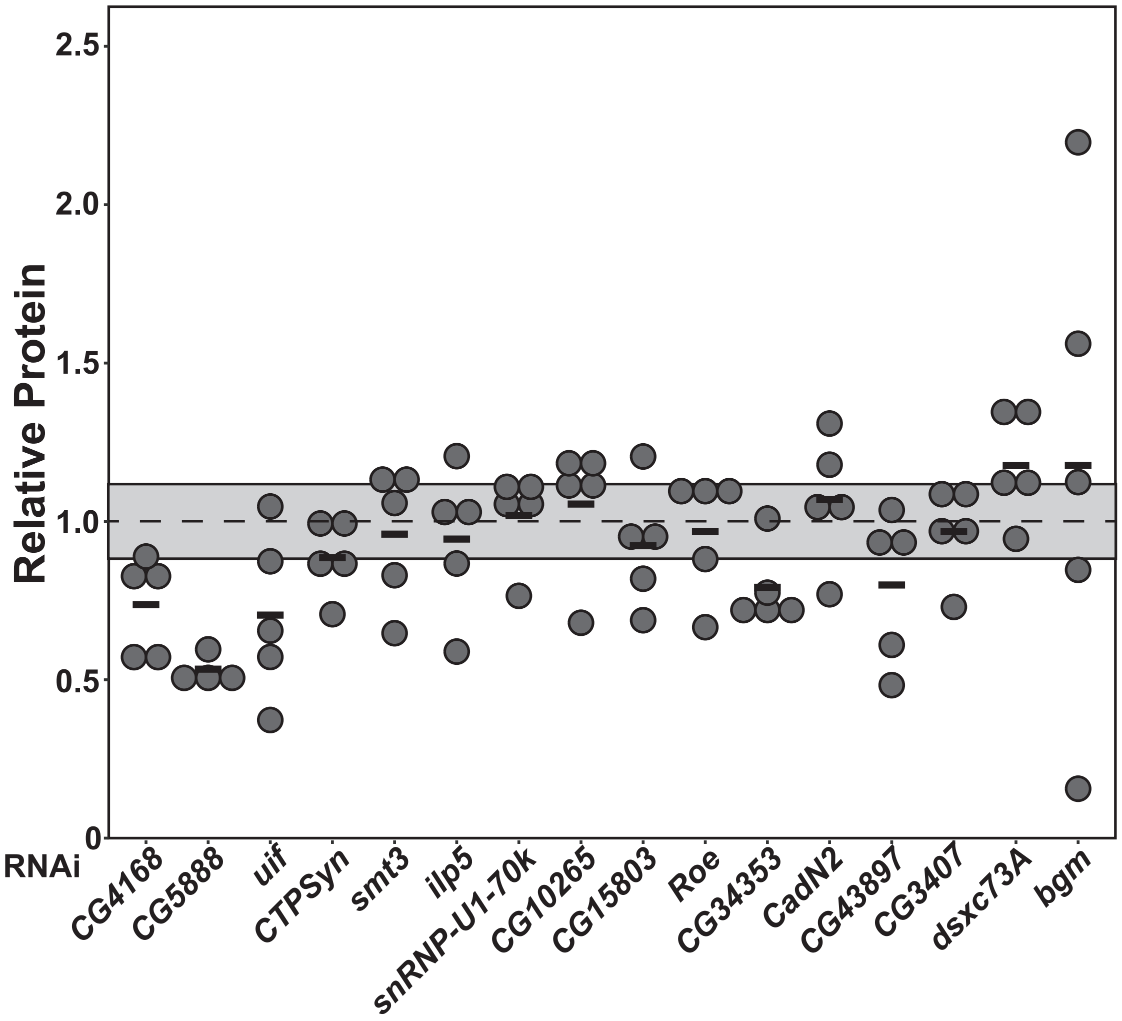


**Supplemental Figure 7. Protein levels are not significantly affected by modifier RNAi**
Protein was measured from the same isolates as glucose (**Figure 4**). Protein level for each sample was normalized to the levels in a genetically matched control line crossed into the *R4-Sirt1* line. Average *R4>Sirt1i* control protein levels after normalization are indicated by a dotted line at 1.0, with standard deviation highlighted by the gray box. Whole fly protein concentration was quantified for N = 4-5 samples per strain, each consisting of 5 flies and individually plotted along the y-axis. Protein levels were not significantly affected by RNAi targeting the indicated candidate modifiers. P-values were calculated using one-way ANOVA followed by Dunnett's multiple testing correction.

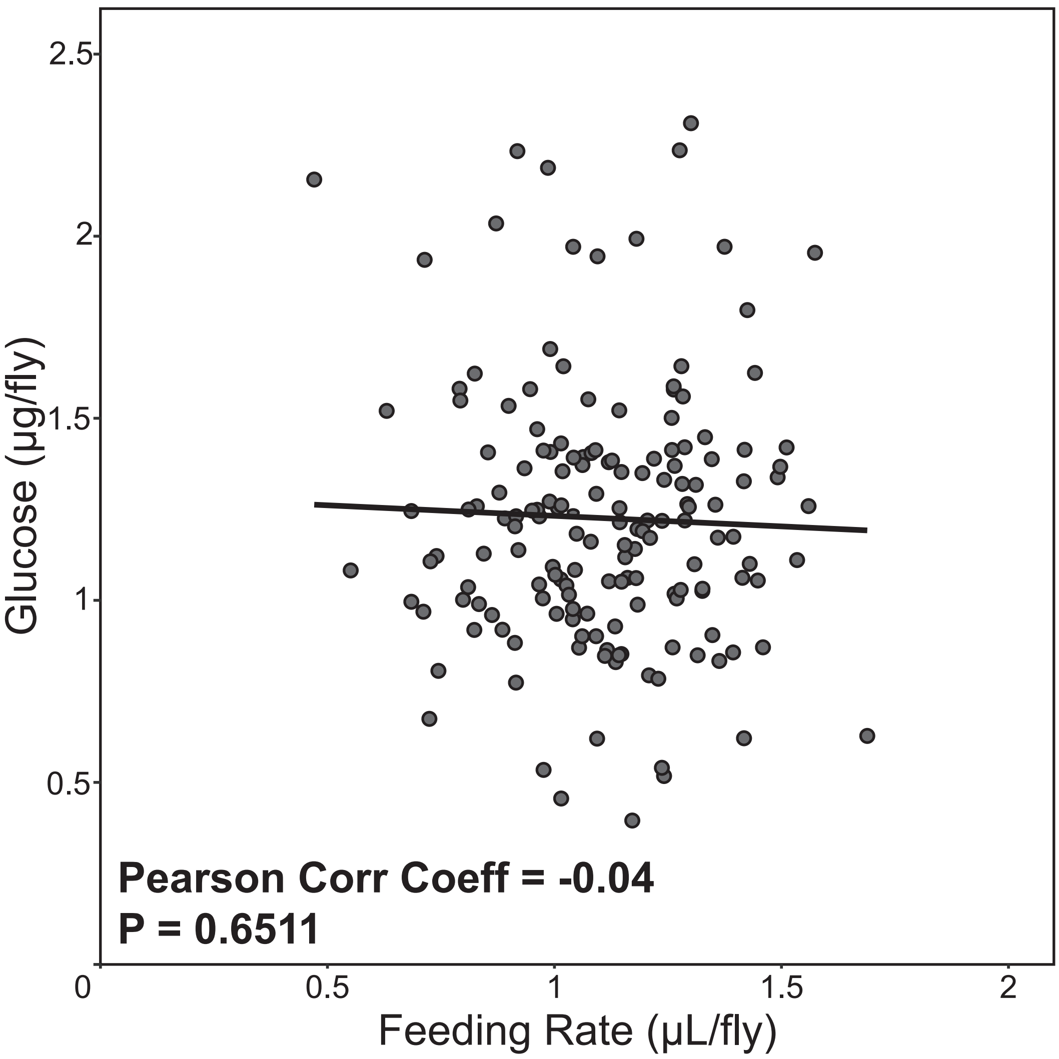


**Supplemental Figure 8. Glucose levels across the DGRP do not correlate with feeding rate.**

Correlation coefficients and p-values were calculated using a Pearson Correlation test. Feeding rate data was obtained from Garlapow *et al.* 2015. Glucose concentrations (µg/fly) across different DGRP strains are not correlated with feeding rate (µL/fly) in the same strains.
